# Antimicrobial resistance determinants in silage

**DOI:** 10.1101/2021.12.12.472282

**Authors:** Sára Ágnes Nagy, Adrienn Gréta Tóth, Márton Papp, Selçuk Kaplan, Norbert Solymosi

## Abstract

Animal products may play a role in developing and spreading antimicrobial resistance in several ways. On the one hand, residues of antibiotics not adequately used in animal farming can enter the human body via food. But resistant bacteria may also be present in animal products, which can transfer the antimicrobial resistance genes (ARG) to the bacteria in the consumer’s body by horizontal gene transfer. As previous studies have shown that fermented foods have a meaningful ARG content, it is indicated that such genes may also be present in silage used as mass feed in the cattle sector. In our study, we aspired to answer what ARGs occur in silage and what mobility characteristics they have? For this purpose, we have analyzed bioinformatically 52 freely available deep sequenced silage samples from shotgun metagenome next-generation sequencing. A total of 17 perfect matched ARGs occurred 55 times in the samples. More than half of these ARGs are mobile because they can be linked to integrative mobile genetic elements, prophages or plasmids. Our results point to a neglected but substantial ARG source in the food chain.

## Introduction

In intensive cattle farming, silage is an essential component of feed. An average dairy cow consumes 25-27 kg of this forage a day, reaching up to a sliage consumption of 12,500 kg per lactation.^1,2^ Silage is most commonly produced of maize or legume plants by anaerobic fermentation. Throughout the fermentation process, fermenting microorganisms, including bacteria, multiply and enrich the silage with beneficial nutrients by the biochemical transformation of its ingredients. If bacteria involved in the process harbour antimicrobial resistance genes (ARGs), the amount of these genes in the silage will increase in parallel with the bacterial counts. Consequently, silage, as a mass feed may continuously supply the gastrointestinal tract of animals with bacteria carrying ARGs. Bacteria entering the digestive system may come into contact with the host microbiota that facilitates the exchange of bacterial genes (e.g. ARGs) by horizontal gene transfer (HGT). HGT may take place as a result of three different processes: conjugation, transduction and transformation. Except for transformation, by which a bacterium can take up any gene from its environment, the routes of HGT require particular active delivery processes. By conjugation, cell-to-cell contact provides the opportunity for a copy of a plasmid to translocate to a recipient bacterium.^3^ Transduction negates the condition of cell-to-cell contact, as in this case, bacteriophages act as a conduit for shuttling genes among bacteria.^4^ The genetic environment of the genes involved in the transfer significantly influences the efficacy of the latter two HGT processes, i.e., the genes’ mobility. The reason why the mobility caharcteristics of ARGs involved in silage are worth taking theinto consideration is the following. If ARGs from silage are transmitted to pathogenic bacteria within an animal’s body, efficacy of antibiotic (AB) treatment may be reduced on the consequent bacterial diseases. In addition, in case of the gut colonization of silage-borne bacteria that carry ARGs, the appearance and enrichment of bacterial ARGs may take place in the animals’ environment after defecation. Decreased efficacy of AB treatments may result in economic loss, and the increased environmental ARG level may have additional veterinary and human health consequences. It proven in former publicationsthat the number of ARGs in fermented dairy products can increase due to the multiplication of fermenting bacteria.^5^ Nevertheless, the description of this phenomenon cannot be found for silage in the literature. Our study aimed to examine the diversity, bacterial relatedness and mobility potential of ARGs deriving from silage. For this purpose, freely available next-generation sequencing (NGS) shotgun metagenome datasets generated by were analyzed by a unified bioinformatics pipeline.

## Materials and Methods

### Data

We searched appropriate datasets in the National Center for Biotechnology Information (NCBI) Sequence Read Archive (SRA) repository. In December 2021, we found only two BioProjects (PRJNA495415, PRJNA764355) sequenced by shotgun metagenomic that had adequate depth for the de novo assembly our study is based on. The median read count (interquartile range, IQR) of the samples was 26.5*×*10^6^ (3.0*×*10^6^) and 34.7*×*10^6^ (1.5*×*10^6^) in datasets PRJNA495415 and PRJNA764355, respectively. There is little metadata available for the samples in the NCBI SRA database only (Table 1). All that can be supposed from the metadata is that the PRJNA495415 samples were taken at different fermentation periods. Samples were taken on days 0, 7, 14, and 28 were classified into groups A, B, C, and D, respectively. Based on the metadata of the PRJNA764355 samples, no such stratification was possible, so all samples were classified in Group E.

**Table 1.**
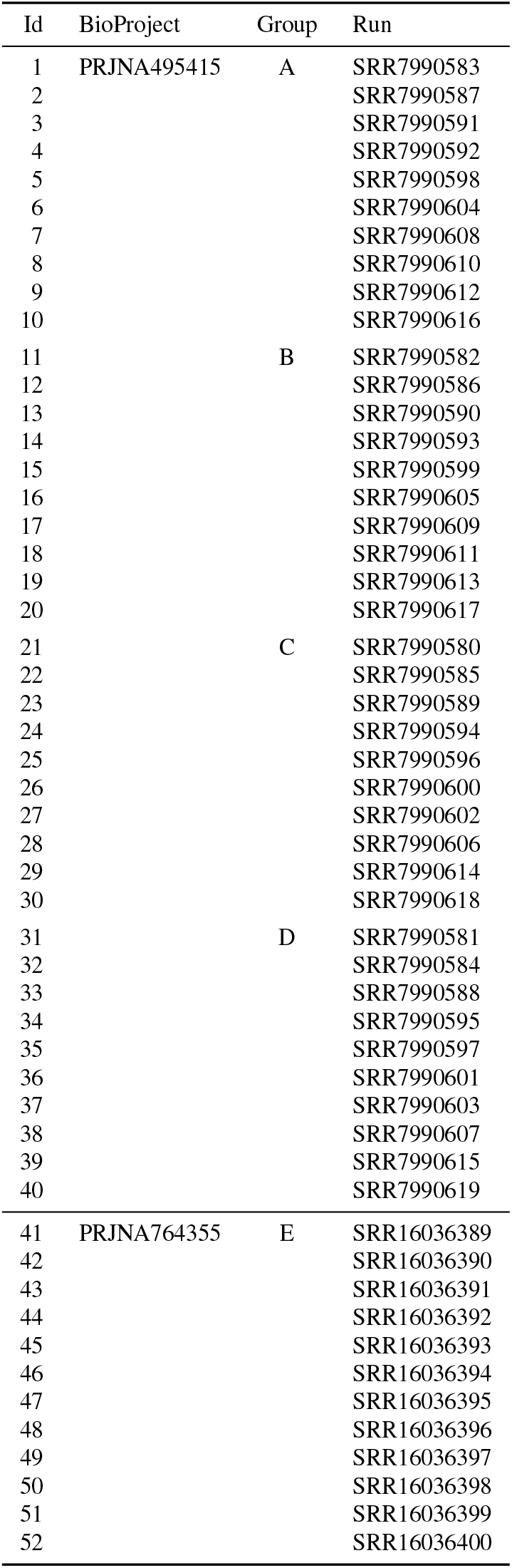
Analyzed samples. The samples of the PRJNA495415 dataset were taken on days 0, 7, 14, and 28 were classified into groups A, B, C, and D, respectively. All samples from BioProject PRJNA764355 are assigned to group E. The run column contains the National Center for Biotechnology Information (NCBI) Sequence Read Archive (SRA) run identifiers.

### Bioinformatic analysis

Quality based filtering and trimming of the raw short reads was performed by TrimGalore (v.0.6.6, https://github.com/FelixKrueger/TrimGalore), setting 20 as a quality threshold. Only reads longer than 50 bp were retained and taxonomically classified using Kraken2 (v2.1.1)^6^ and a database created (24/03/2021) from the NCBI RefSeq complete archaeal, bacterial, viral and plant genomes. For this taxon assignment the -confidence 0.5 parameter was used to obtain more precise species level hits. The taxon classification data was managed in R^7^ using functions of the packages phyloseq^8^ and microbiome.^9^ The preprocessed reads were assembled to contigs by MEGAHIT (v1.2.9)^10^ using default settings. The contigs were also classified taxonomically by Kraken2 with the same database as above. From the contigs all possible open reading frames (ORFs) were gathered by Prodigal (v2.6.3)^11^. The protein translated ORFs were aligned to the ARG sequences of the Comprehensive Antibiotic Resistance Database (CARD, v.3.1.3)^12,13^ by Resistance Gene Identifier (RGI, v5.2.0) with Diamond^14^. ORFs having a perfect match against the CARD database were kept only for further analysis. The integrative mobile genetic element (iMGE) content of the ARG harbouring contigs was analyzed by MobileElementFinder (v1.0.3) and its database (v1.0.2).^15^ Following the distance concept of Johansson et al.^15^ for each bacterial species, those with a distance threshold defined within iMGEs and ARGs were considered associated. In the MobileElementFinder database (v1.0.2) for *Enterobacter hormaechei*, the longest composite transposon (cTn) was the *Tn3000*. In the case of this species, its length (11,823 bp) was taken as the cut-off value. For *Enterococcus faecium*, this threshold was the length of the *Tn6246* transposon, 5,147 bp. As the database neither contains species-level, nor genus-level cTn data for *Bacillus, Lactiplantibacillus* and *Lacticaseibacillus* species, a general cut-off value was chosen for the contigs of these species. This value was declared as the median of the longest cTns per species in the database (10,098 bp). The plasmid origin probability of the contigs was estimated by PlasFlow (v.1.1)^16^ The phage content of the assembled contigs was prediced by VirSorter2 (v2.2.3)^17^. The findings were filtered for dsDNAphages and ssDNAs. All data management procedures, analyses and plottings were performed in R environment (v4.1.0).^7^

## Results

Based on the taxon classification performed on a database containing plant complete reference genomes, the dominant plants in the silage belong to the *Medicago* genus, most likely alfalfa (*M. sativa*). Further analysis results of the shotgun metagenomic sequenced 52 samples (Table 1) is summarised in the following sections. After presenting the bacteriome and the identified AGRs (resistome), predictions regarding the mobility potential of ARGs were also resumed based on genetic characteristics that may play a significant role in HGT.

### Bacteriome

By taxon classification, the number of reads aligning to bacterial genomes varied by samples (median: 20.6*×*10^6^, IQR: 2.9*×*10^6^). The relative abundances of genera that achieved more than 1% of the bacterial hits in any of the samples is shown in Figure 1. The dominant bacteria genera (with mean abundance) in descending order were *Weissella* (45.7%), *Pantoea* (18.5%), *Levilactobacillus* (13.5%), *Pediococcus* (6.7%), *Lactiplantibacillus* (6.3%), *Companilactobacillus* (1.7%), *Lacticaseibacillus* (1.3%), *Enterococcus* (1.2%), *Lactococcus* (1%), *Kosakonia* (0.8%), *Staphylococcus* (0.6%), *Enterobacter* (0.5%), *Latilactobacillus* (0.5%), *Bacillus* (0.4%), *Limosilactobacillus* (0.4%), *Pseudomonas* (0.4%), *Leclercia* (0.2%), *Mammaliicoccus* (0.2%), *Agrobacterium* (0.1%).

**Figure 1.**
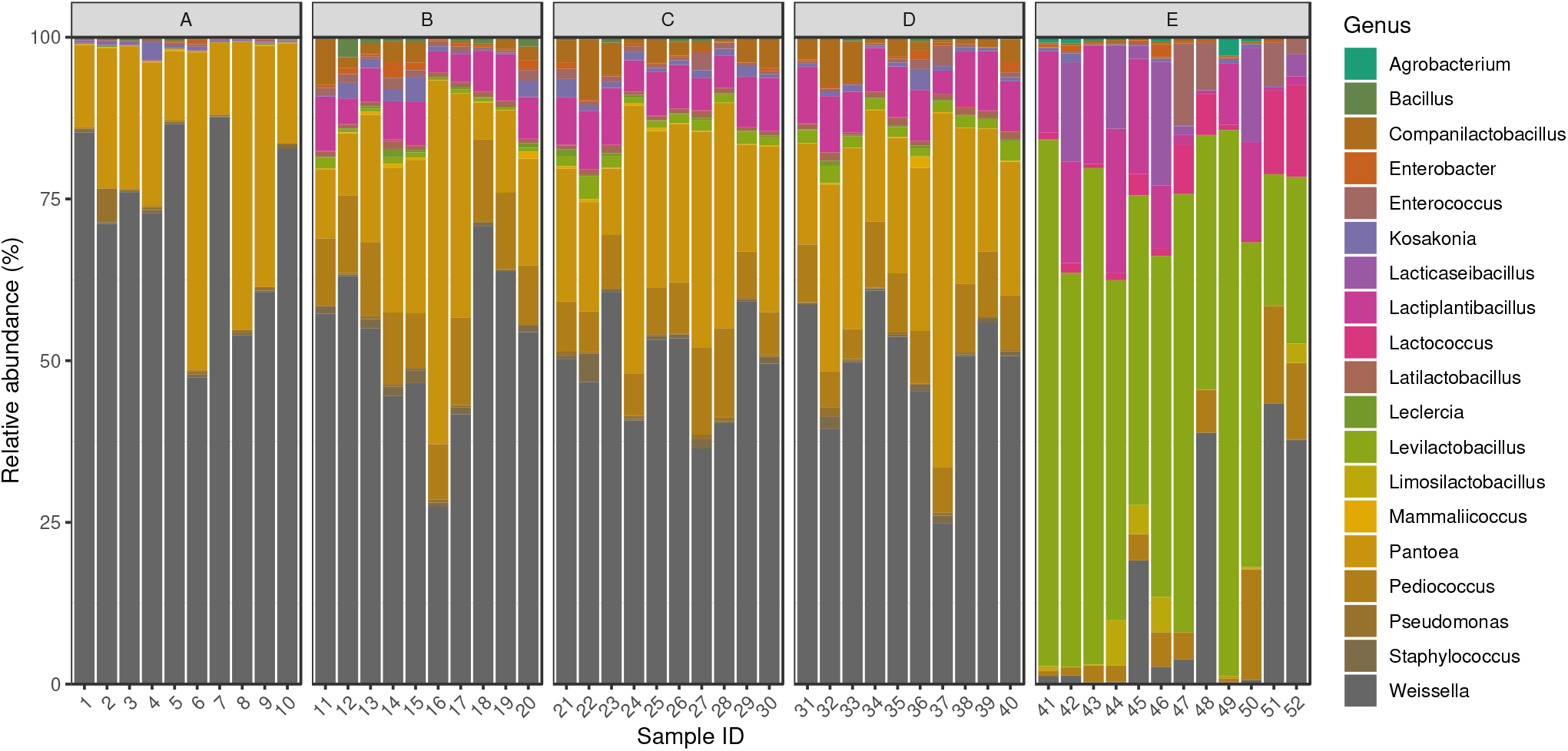
Silage core bacteriome. The relative abundances of genera that achieved more than 1% of the bacterial hits in any of the samples. The elements of the PRJNA495415 dataset were taken on days 0, 7, 14, and 28 were classified into groups A, B, C, and D, respectively. All items from BioProject PRJNA764355 are assigned to group E.

### Resistome

The median length of the filtered contigs harbouring ARGs constructed by de novo assembly was 4,137 bp (IQR: 2,911). The number of ARGs found on the contigs ranged from 1 to 2. From the analyzed 52 samples in 20 occured 55 times the identifed 17 various ARGs: *aadA2, ANT(6)-Ia, ANT(9)-Ia, APH(3’)-IIa, APH(3’)-IIIa, dfrG, Erm(44)v, lmrD, lsaE, poxtA, qacEdelta1, QnrD1, QnrS1, sul1, sul2, tet(K), vatE* (Figure 2).

**Figure 2.**
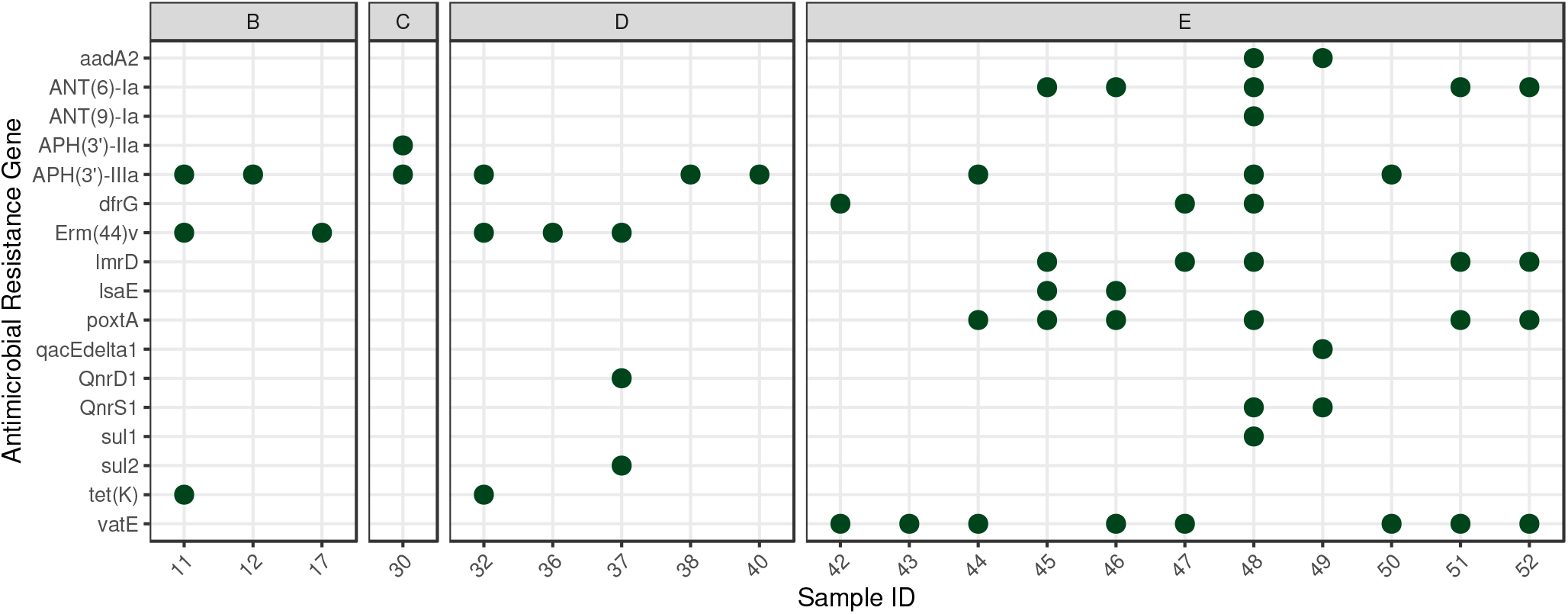
Identifed antimicrobial resistance genes (ARGs) by samples. The perfect matched ARGs were plotted by samples. The elements of the PRJNA495415 dataset were taken on days 0, 7, 14, and 28 were classified into groups A, B, C, and D, respectively. All samples from BioProject PRJNA764355 are assigned to group E.

The resistance mechanism of identified ARGs was in descending order of frequency the antibiotic inactivation (47.3%), antibiotic target protection (20%), antibiotic efflux (14.5%), antibiotic target alteration (9.1%), antibiotic target replacement (9.1%).

The identified ARGs associated with bacteria by species are as follows. *Acinetobacter baumannii*: *aadA2, qacEdelta1*; *Amylolactobacillus amylophilus*: *ANT(6)-Ia*; *Bacillus subtilis*: *APH(3’)-IIa*; *Cronobacter* sp. JZ38: *QnrS1*; *Enterobacter hormaechei*: *aadA2, sul1*; *Enterococcus faecalis*: *APH(3’)-IIIa*; *E. faecium*: *poxtA*; *Escherichia coli*: *sul2*; *Gracilibacillus* sp. SCU50: *dfrG*; *Lacticaseibacillus manihotivoran*s: *ANT(6)-Ia*; *L. paracasei*: *poxtA*; *Lactiplantibacillus plantarum*: *APH(3’)-IIIa, poxtA, vatE*; *Lactococcus lactis*: *lmrD*; *Levilactobacillus brevis*: *poxtA*; *Ligilactobacillus acidipiscis*: *ANT(9)-Ia*; *Providencia rettgeri*: *QnrD1*; *Staphylococcus aureus*: *ANT(6)-Ia, tet(K)*; *S. carnosus*: *Erm(44)v*; *S. pseudoxylosus*: *Erm(44)v*; *S. saprophyticus*: *Erm(44)v*; *Streptococcus suis*: *lsaE*; *Tetragenococcus halophilus*: *APH(3’)-IIIa*; *Weissella paramesenteroides*: *ANT(6)-Ia*.

### Mobilome

We found a total of 53 ARGs that can be assumed to be mobile. Ten of these ARGs are linked to integrative mobile genetic elements (iMGE). Further, 2 ARGs were detected in prophages and 41 on plasmids. The frequencies of ARGs associated with iMGEs, phages and plasmids by bacteria species of origin are summarized in Figure 3.

**Figure 3.**
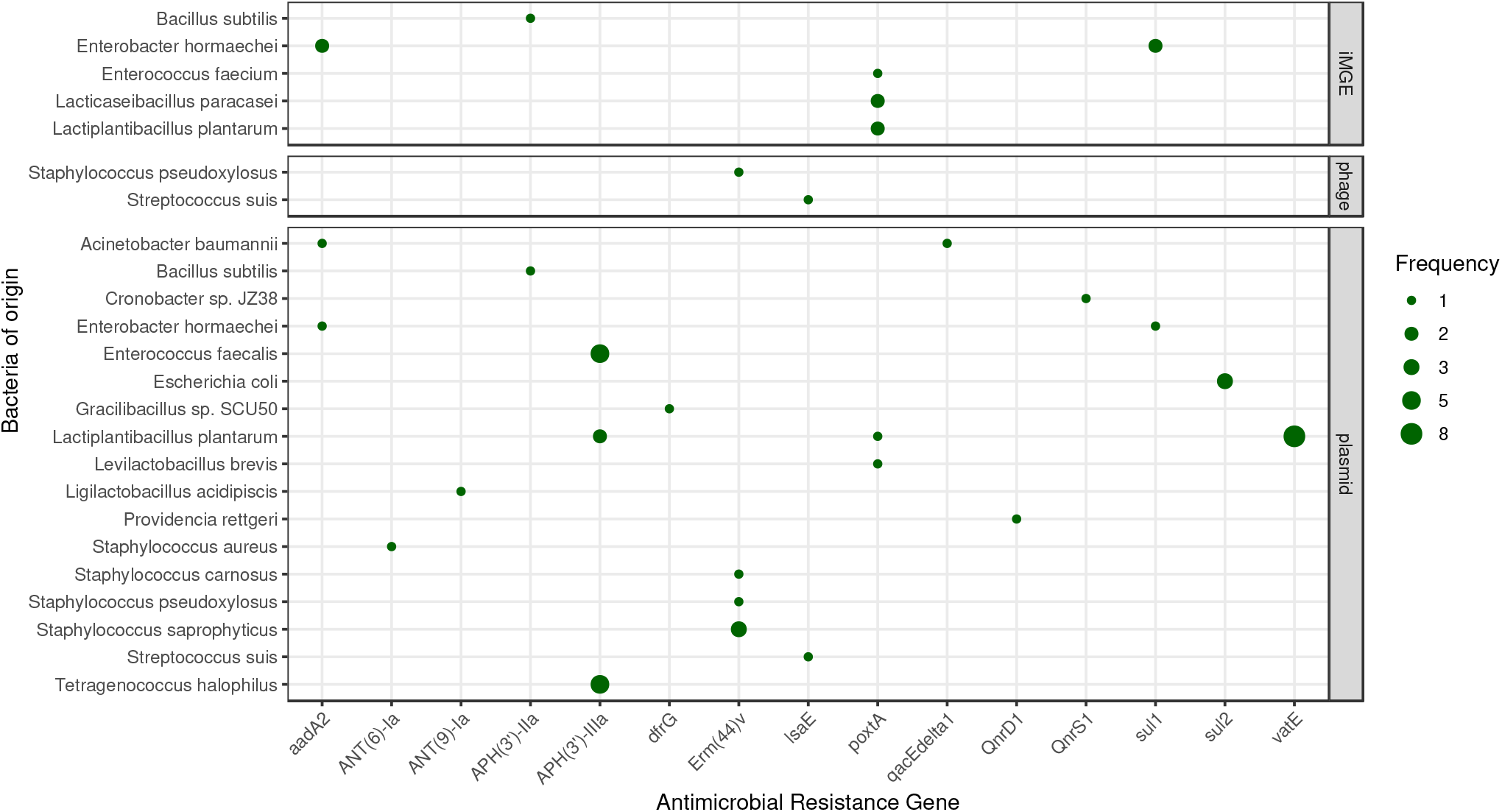
Mobile antimicrobial resistance gene frequency by bacteria of origin. The size of the dots indicates the occurrence frequency of the given gene flanked by iMGE, positioned in plasmid or phage.

Following the distance method proposed by Johansson et al. (2021)^15^ integrated mobile genetic element associated ARGs were detected in five samples (30, 45, 46, 48, 52) and five species (*Bacillus subtilis, Enterobacter hormaechei, Enterococcus faecium, Lacticaseibacillus paracasei, Lactiplantibacillus plantarum*). The *Bacillus subtilis* carried *APH(3’)-IIa* in sample 30, the *poxtA* of *Enterococcus faecium* in sample 45, *Lactiplantibacillus plantarum* in sample 52, and *Lacticaseibacillus paracasei* in sample 46 was detected as iMGE linked gene. The *Enterobacter hormaechei* harboured *sul1* and *aadA2* genes co-existed with integrated mobile elements in sample 48. Two prophage-linked ARGs were identified, the contig harbouring *Erm(44)v* classified to *Staphylococcus pseudoxylosus* by VirSorter2 was found to be of dsDNA phage origin. While the contig of *lsaE* from *Streptococcus suis* was predicted as ssDNA derived. The previous one was detected in sample 37, the latter in sample 45. In 19 samples (Nr. 11, 12, 17, 30, 32, 36, 37, 40, 42, 43, 44, 45, 46, 47, 48, 49, 50, 51, 52) were contigs harbouring ARG predicted plasmid of origin.

## 1 Discussion

Our study identified numerous perfect matched ARGs in Medicago silage samples. All but group A had at least one sample containing one or more ARGs. Among the PRJNA495415 Bioproject samples, the highest number of ARGs were found in group D. The interpretation opportunities of these finding is limited as we do not know much about the samples. However, the results suggest that bacteria that carry ARG may be multiplied during the fermentation process. Interestingly, all but one of the PRJNA764355 bioproject samples contained ARG. Due to the lack of metadata, it is hard to find any reason for this high ARG level. However, one possible cause might be that the PRJNA764355 samples were sequenced by approximately 1.3 times more reads than the PRJNA495415 samples. It is known from previous studies that with deeper sequencing, more complete genes can be generated by de novo assembly.^5,18^

The carrier bacteria predicted for the identified ARGs can be classified according to their presence in silage. In the literature, one can find the following bacteria as a characteristic feature of silage: *Bacillus subtilis*^19^ *Enterococcus faecium*^20^ *Escherichia coli*^21^, *Lactiplantibacillus plantarum*^22^, *Lactococcus lactis*^23^, *Levilactobacillus brevis*^24–26^, *Ligilactobacillus acidipiscis*^23^ *Weissella paramesenteroides*^27^ The genera of these species dominate the bacteriome of the samples. The identified *Cronobacter sp. JZ38*^28^ may be of plant origin. However, it can be assumed that other species may be present as contaminants of the silage: *Amylolactobacillus amylophilus, Enterobacter hormaechei, Enterococcus faecalis, Gracilibacillus* sp. SCU50, *Lacticaseibacillus manihotivorans, Lacticaseibacillus paracasei, Providencia rettgeri, Staphylococcus aureus, Staphylococcus carnosus, Staphylococcus pseudoxylosus, Staphylococcus saprophyticus, Streptococcus suis, Tetragenococcus halophilus*. Nevertheless, some of these bacteria are members of the *Lactobacillaceae* family, the *Leuconostoc* or *Enterobacter* genera. Which groups’ numerous species are typical of fermented food and feed components.

The following was found in the literature regarding the co-occurrence of the ARGs identified in our study and the bacteria carrying them. *AadA2* encoding an aminoglycoside nucleotidyltransferase and *QacEdelta1*, a resistance gene conferring resistance to antiseptics have both been described in *Acinetobacter baumanni* in former publications^29,30^. *ANT(6)-Ia*, that is an aminoglycoside nucleotidyltransferase gene, appears in many species, including *Lactobacillus* spp.^31^. Its species-specific association with *Amylolactobacillus amylophilus* has not been described in any former publications. *APH(3’)-IIa*, an aminoglycoside phosphotransferase^13^, to our knowledge, has not detected in *Bacillus subtilis* up till now. *QnrS1* encoding a quinolone resistance protein was originally identified in *Shigella flexneri*^32^. In line with our results, this gene has recently been mentioned to appear in *Cronobacter* spp. in a case report^33^. *Enterobacter hormaechei* deriving *aadA2* and *sul1*, a sulfonamide resistant dihydropteroate synthase gene that is described to appear in Gram-negative bacteria^13^ have been reported to appear in the genom of *Enterobacter* spp. and *Enterobacter hormaechei*, respectively in former publications as well^34,35^. Within the *Enterococcus* genus, two perfect ARG matches were identified, namely *APH(3’)-IIIa* in *Enterococcus faecalis* and *poxtA* in *E. faecium. APH(3’)-IIIa* is an aminoglycoside phosphotransferase that normally appears in *Staphylococcus aureus*^13^ and *Enterococcus* spp.woegerbauer2015involvement, while *poxtA* is a gene encoding an ABC-F subfamily (ATP-binding cassette-F) protein that facilitates resistance to tetracycline, phenicol, and oxazolidinone via modification of the bacterial ribosome. First detection of *poxtA* took place in a methicillin-resistant *Staphylococcus aureus* strain^13^, followed by other bacterial species, including *Enterococcus faecium*^36^. *Sul2*, a sulfonamide resistant dihydropteroate synthase of Gram-negative bacteria is commonly described in *Escherichia coli*^13,37^. *dfrG* is a plasmid-encoded dihydrofolate reductase^13^ that, to our knowledge, has not been described in *Gracibacillus* spp. up till now, but has already appeared in *Bacillaceae* family^38^. *ANT(6)-Ia*, an aminoglycoside nucleotidyltransferase gene appears in many species, including *Lactobacillus* spp.^31^. Its species-specific association with *Lacticaseibacillus manihotivorans* has not been described in any publications. *PoxtA* that was detected in *Lacticaseibacillus paracasei, Lactiplantibacillus plantarum* and *Levilactobacillus brevis* in the silage samples, has been described to appear in *Lactobacillaceae*, namely *Lactobacillus acidophilus*, but not in these very species^39^. Another species that was detected harboring *APH(3’)-IIIa* in the silage samples was *Lactiplantibacillus plantarum*. This finding is in line with the ARG-species match results mentioned in former publications^40^. Furthermore, *L. plantarum* was also associated with *vatE* that encodes an acetyltransferase conferring resistance against streptogramins^13^. *VatE* was originally found in *Enterococcus faecium*^13^ and has since then been identified in *Lactobacillaceae*^41^, but not specifically in *L. plantarum. Ligilactobacillus acidipiscis ANT(9)-Ia*, an aminoglycoside nucleotidyltransferase gene^13^ was associated with this genus for first within this study. *QnrD1*, a gene encoding a quinolone resistance protein that is normally detected in *Salmonella enterica*^13^, has already been found in *Providencia* spp.^42^ and was attached to *Providencia rettgeri* in our study as well. *Staphylococcus aureus* could have been associated with two ARGs, *ANT(6)-Ia* and *tetK* encoding a tetracycline efflux protein, that are both common findings in *Staphylococcus* spp.^43,44^. Although, *Erm(44)v* was first detected in the *Staphylococcus saprophyticus*^45^, no literature could be found about the appearance of this gene in *Staphylococcus carnosus*, in *S. nepalensis* or in *S. pseudoxylosus* species. *lsaE* encoding another ABC-F subfamily protein conferring resistance to pleuromutilin, lincosamide, and streptogramin A is a common finding in *Streptococcus* spp.^46^ and has also been associated with *S. suis* in previous publications^47^. Besides the bacterial species mentioned above, *APH(3’)-IIIa* was also detected in *Tetragenococcus halophilus*. This ARG is often appears in *Enterococcaceae*^13^ but has not yet been written down in this species. Furthermore, to our knowledge, *Weissella paramesenteroides* associated *ANT(6)-Ia* has been detected for first within this study.

The ARGs belonging to the genome of *Acinetobacter baumannii* may play a role in the appearance of resistance against acridine dye, aminoglycoside, disinfecting agents and intercalating dyes; *Amylolactobacillus amylophilus*: aminoglycoside; *Bacillus subtilis*: aminoglycoside; *Cronobacter* sp. JZ38: fluoroquinolone; *Enterobacter hormaechei*: aminoglycoside, sulfonamide; *Enterococcus faecalis*: aminoglycoside; *E. faecium*: lincosamide, macrolide, oxazolidinone, phenicol, pleuromutilin, streptogramin, tetracycline; *Escherichia coli*: sulfonamide; *Gracilibacillus* sp. SCU50: diaminopyrimidine; *Lacticaseibacillus manihotivorans*: aminoglycoside; *L. paracasei*: lincosamide, macrolide, oxazolidinone, phenicol, pleuromutilin, streptogramin, tetracycline; *Lactiplantibacillus plantarum*: aminoglycoside, lincosamide, macrolide, oxazolidinone, phenicol, pleuromutilin, streptogramin, tetracycline; *Lactococcus lactis*: lincosamide; *Levilactobacillus brevis*: lincosamide, macrolide, oxazolidinone, phenicol, pleuromutilin, streptogramin, tetracycline; *Ligilactobacillus acidipiscis*: aminoglycoside; *Providencia rettgeri*: fluoroquinolone; *Staphylococcus aureus*: aminoglycoside, tetracycline; *S. carnosus*: lincosamide, macrolide, streptogramin; *S. pseudoxylosus*: lincosamide, macrolide, streptogramin; *S. saprophyticus*: lincosamide, macrolide, streptogramin; *Streptococcus suis*: lincosamide, macrolide, oxazolidinone, phenicol, pleuromutilin, streptogramin, tetracycline; *Tetragenococcus halophilus*: aminoglycoside; *Weissella paramesenteroides*: aminoglycoside.

Throughout our study, several ARGs were predicted to be co-occurring with genetic attributes facilitating mobility. The bioinformatic analysis of the mobility characteristics relied upon the identification of three major mobility determination groups, namely iMGEs, phages and plasmids. We found APH(3’)-IIa linked to integrated mobile genetic element in Bacillus subtilis, in the literature can be find similar for Escherichia coli.^13^ While in the literature the resistance genes of aadA2 and sul1 on plasmids in Enterobacter hormaechei were formerly described^48^, we found them enisled by iMGEs. Our findings on iMGE flanked poxtA in Enterococcus faecium is in line with the current literature.^49^ We found the same co-occurence in Lacticaseibacillus paracasei, but no similar in the literature. Within prophages there were found the gene Erm(44)v and lsaE, in Staphylococcus pseudoxylosus and Streptococcus suis, respectively. While similar like the first linkage can be found in the literature^50^, for the latter one we found only not mobile report^47^. All the other mobile ARGs were detected on contigs predicted as plasmid originated. In the case of aadA2 and qacEdelta1 in Acinetobacter baumannii^13^; aadA2 and sul1 in Enterobacter hormaechei^13^; APH(3’)-IIIa in Enterococcus faecalis^13^; sul2 in Escherichia coli^13^; APH(3’)-IIIa in Lactiplantibacillus plantarum^51^; QnrD1 in Providencia rettgeri^13^ ANT(6)-Ia in Staphylococcus aureus^13^ and Erm(44)v in Staphylococcus saprophyticus^50^ the placement on plasmid is can be found in the literature. We have no data plasmid co-occurence of APH(3’)-IIa in Bacillus subtilis, dfrG in Gracilibacillus sp. SCU50, ANT(9)-Ia in Ligilactobacillus acidipiscis, lsaE in Streptococcus suis, QnrS1 in Cronobacter sp. JZ38, Erm(44)v in Staphylococcus carnosus and S. pseudoxylosus. Hao et al. described poxtA gene in a multiresistance plasmid with mobile elements in Enterococcus fecalis. This gene found in a number of Gram-positive bacteria, including enterococci as well, but it is not found in Lactiplantibacillus plantarum nor Levilactobacillus brevis.^52^ Previous findings confirm vat(E) occures on plasmids.^53^ In spite of its frequent presence in enterococci^54^ there is no evidence of the presence in L. plantarum. We found that the APH(3’)-IIIa gene of Tetragenococcus halophilus encoded by plasmid which is consistent with what has been described in previously, APH(3)IIIa was found on high molecular weight plasmids and chromosomes of the enterococcal species^55^. Although the description of the APH(3’)-IIIa gene in T. halophilus is not found.

The results described above indicate that the silage contains ARGs in some bacteria responsible for its formation, and some ARGs are mobile as well. These mobile ARGs entering the animal can intrude in saprophytic and pathogenic bacteria by horizontal gene transfer. From the farm animal body,^56–58^ the ARG harbouring bacteria can be distributed by products intended for human consumption.^59^ Finally, the ARG transferred by this was into the human body might decrease the effectiveness of antibiotic therapy. However, to get a picture of the whole process, many points still need to be examined and clarified. It would be essential to analyze the colonization success of the ARG harbouring bacteria entering the body from the silage, the extent of the transfer of the ARGs they carry to the bacteria living in the gastrointestinal tract. The silage involved in the study is of Medicago origin, and our results are based on data from only two Chinese projects. So, it would also be necessary to investigate the ARG content of other alfalfa and corn silages.

Antimicrobial resistance is an emerging threat to public health worldwide. This concern affects not only the healthcare sector but also agriculture. The usage of antibiotics in livestock exceeds the rate of human applications^60^. Consequently, the antibiotics used in food animal medicine are considered a grave risk to humans through the food chain^61^. The effects of the veterinary administered antimicrobials on the food chain are well published, whereas the presence of the ARGs in dairy cattle nutrition is underrepresented in the literature. Still, the microbial mass contained in fermented feeds could play an essential role in the ARGs shifting through the food chain.

## Acknowledgements

The authors would like to thank the providers of BioProject PRJNA495415. The research was supported by the European Union’s Horizon 2020 research and innovation program under Grant Agreement No. 874735 (VEO).

## Author contributions statement

NS takes responsibility for the integrity of the data and the accuracy of the data analysis. AGT, MP, NS and SÁN conceived the concept of the study. AGT, MP, NS and SÁN participated in the bioinformatic analysis. AGT, KS, MP, NS and SÁN participated in the drafting of the manuscript. AGT, KS, MP, NS and SÁN carried out the critical revision of the manuscript for important intellectual content. All authors read and approved the final manuscript.

